# Altered Bile Acid Profile in Mild Cognitive Impairment and Alzheimer’s Disease: Relationship to Neuroimaging and CSF Biomarkers

**DOI:** 10.1101/284141

**Authors:** Kwangsik Nho, Alexandra Kueider-Paisley, Siamak MahmoudianDehkordi, Matthias Arnold, Shannon L. Risacher, Gregory Louie, Colette Blach, Rebecca Baillie, Xianlin Han, Gabi Kastenmüller, Wei Jia, Guoxiang Xie, Shahzad Ahmad, Thomas Hankemeier, Cornelia M. van Duijn, John Q. Trojanowski, Leslie M. Shaw, Michael W. Weiner, P. Murali Doraiswamy, Andrew J. Saykin, Rima Kaddurah-Daouk, for the Alzheimer’s Disease Neuroimaging Initiative and the Alzheimer Disease Metabolomics Consortium

**Affiliations:** Department of Radiology and Imaging Sciences, Center for Computational Biology and Bioinformatics, and the Indiana Alzheimer Disease Center, Indiana University School of Medicine, Indianapolis, IN, USA; Department of Psychiatry and Behavioral Sciences, Duke University, Durham, NC, USA; Institute of Bioinformatics and Systems Biology, Helmholtz Zentrum München, German Research Center for Environmental Health, Neuherberg, Germany; German Center for Diabetes Research (DZD), Neuherberg, Germany; Duke Molecular Physiology Institute, Duke University, Durham, NC, USA; Rosa & Co LLC, San Carlos, CA, USA; University of Texas Health Science Center at San Antonio, San Antonio, TX, USA; University of Hawaii Cancer Center, Honolulu, Hawaii, USA; Erasmus Medical Centre, Rotterdam, the Netherlands; Division of Analytical Biosciences, Leiden Academic Centre for Drug Research, Leiden University, P.O. Box 9502, 2300 RA Leiden, the Netherlands; Department of Pathology & Laboratory Medicine, University of Pennsylvania, Philadelphia, PA, USA; Center for Imaging of Neurodegenerative Diseases, Department of Radiology, San Francisco VA Medical Center/University of California San Francisco, San Francisco, CA; Duke Institute of Brain Sciences, Duke University, Durham, NC, USA; Department of Medicine, Duke University, Durham, NC, USA

**Keywords:** Metabolomics, bile acid, Alzheimer’s disease, amyloid-β, CSF biomarkers, brain glucose metabolism, PET, MRI, gut liver brain axis

## Abstract

**Introduction:** Bile acids (BAs) are the end products of cholesterol metabolism produced by human and gut microbiome co-metabolism. Recent evidence suggests gut microbiota influence pathological features of Alzheimer’s disease (AD) including neuroinflammation and amyloid-β deposition.

**Method:** Serum levels of 20 primary and secondary BA metabolites from the AD Neuroimaging Initiative (n=1562) were measured using targeted metabolomic profiling. We assessed the association of BAs with the “A/T/N” (Amyloid, Tau and Neurodegeneration) biomarkers for AD: CSF biomarkers, atrophy (MRI), and brain glucose metabolism ([^18^F]FDG-PET).

**Results:** Of 23 BA and relevant calculated ratios, three BA signatures were associated with CSF Aβ1-42 (“A”) and three with CSF p-tau181 (“T”) (corrected *p*<0.05). Furthermore, three, twelve, and fourteen BA signatures were associated with CSF t-tau, glucose metabolism, and atrophy (“N”), respectively (corrected *p*<0.05).

**Conclusion:** This is the first study to show serum-based BA metabolites are associated with “A/T/N” AD biomarkers, providing further support for a role of BA pathways in AD pathophysiology. Prospective clinical observations and validation in model systems are needed to assess causality and specific mechanisms underlying this association.

## 1. Introduction

Several metabolic perturbations have been noted in Alzheimer’s disease (AD) including failures associated with cholesterol metabolism[1-3] which has been associated with AD in multiple lines of research including physiological and epidemiological studies.[3-5] Cholesterol is synthesized in liver and its clearance involves bile acid (BA) production by gut microbiome and human co-metabolism. Changes in microbial gut populations can profoundly alter BA profiles and signaling. Bile acids appear to play a role in the central nervous system.[6, 7] Recent work suggests microbial disturbances linked to BA profiles are implicated in neurodegenerative disorders.[8-13] The gut microbiota are involved in immune, neuroendocrine, and neural pathways[10, 11, 14-18], have been shown to regulate microglial maturation and function, and may contribute to AD.[18, 19]

Peripheral metabolic changes may influence central changes through the liver and gut-brain axis that includes commensal and pathogenic bacteria, through its interactions with the vagus nerve, changes in central nervous system functioning, the immune system[20, 21], and hippocampal neurogenesis.[22] These signals are crucial for the regulation of energy, glucose homeostasis and inflammation.[23] The gut-brain biochemical axis of communication is just starting to be elucidated. Circulating BAs seem to provide an important mechanism for communication between the gut and the brain, and their alterations reflects gut dysbiosis. Previous studies suggest that BAs are altered in mild cognitive impairment (MCI) and AD[24], and in the preceding paper, we showed that increased levels of secondary cytotoxic BAs and their ratios to primary BAs were associated with AD and poor cognition. This supported the hypothesis that circulating BAs may contribute to AD pathogenesis. Research in AD animal models suggests a role for the gut microbiome in the development of amyloid-β pathology[25]. However, little work has been done in humans to link peripheral metabolic changes in cholesterol to central biomarkers related to AD including amyloid-β and tau accumulation, brain glucose metabolism, and structural atrophy. Therefore, we analyzed serum BA metabolites and their ratios from older adults with early stage AD or who were at risk for AD from the Alzheimer’s Disease Neuroimaging Initiative (ADNI) cohort.

We hypothesized that serum BA levels and their relevant ratios would associate with biomarkers of AD pathophysiology including neuroimaging (MRI and PET) and cerebrospinal fluid (CSF). The AD biomarkers were selected and defined consistent with the emergent NIA-Alzheimer’s Association Research Framework (“A/T/N”) for AD biomarkers that defines three general groups of biomarkers based on the nature of pathologic process that each measures (https://www.alz.org/aaic/_downloads/nia-aa-draft-11-27-2017.pdf).[26, 27] Biomarkers of amyloid-β plaque (“A”) are CSF Aβ1-42 and cortical amyloid-β accumulation measured by Florbetapir PET, biomarkers of fibrillary tau (“T”) are CSF phosphorylated tau (p-tau) and cortical tau deposition measured by tau PET, and biomarkers of neurodegeneration or neuronal injury (“N”) are atrophy on MRI, glucose metabolism on FDG PET, and CSF total tau (t-tau).

## 2. Methods

### 2.1. Study cohort

Serum samples and data analyzed in the present report were obtained from ADNI. The initial phase (ADNI-1) was launched in 2003 to test whether serial magnetic resonance imaging (MRI), position emission tomography (PET), other biological markers, and clinical and neuropsychological assessment could be combined to measure the progression of MCI and early AD. ADNI-1 was extended to subsequent phases (ADNI-GO, ADNI-2, and ADNI-3) for follow-up for existing participants and additional new enrollments. Inclusion and exclusion criteria, clinical and neuroimaging protocols, and other information about ADNI can be found at www.adni-info.org.[28, 29] Demographic information, raw neuroimaging scan data, *APOE*, neuropsychological test scores, and clinical information are available and were downloaded from the ADNI data repository (www.loni.usc.edu/ADNI/). Written informed consent was obtained at the time of enrollment that included permission for analysis and data sharing and consent forms were approved by each participating sites’ Institutional Review Board (IRB).

### 2.2. Quality control procedures of serum bile acid profiles

Targeted metabolomics profiling was performed to identify and quantify concentrations of 20 BAs from serum samples using <u>Biocrates^®^ Bile Acids Kit</u> as described in detail [previous paper]. In brief, morning serum samples from the baseline visit were collected and aliquoted as described in the ADNI standard operating procedures, with only fasting samples included in this study.[28] BA quantification was performed by liquid chromatography tandem mass spectrometry. Metabolites with >40% of measurements below the lower limit of detection (<LOD) were excluded. To assess the precision of the measured analytes, a set of blinded analytical replicates (24 pairs in ADNI-1 and 15 triples in ADNI-GO/2) were supplied by ADNI. Unblinded metabolite profiles went through further quality control (QC) checks. The preprocessed dataset included 15 BAs (5 BAs did not pass QC criteria) and 8 ratios. These selected ratios reflect enzymatic dysfunctions in liver and changes in gut microbiome metabolism (**Fig 1b**). The preprocessed BA values obtained from the QC step were adjusted for the effect of medication use (at baseline) on BA levels (see Toledo et al. 2017 for adjustment description details).

**Fig 1.**
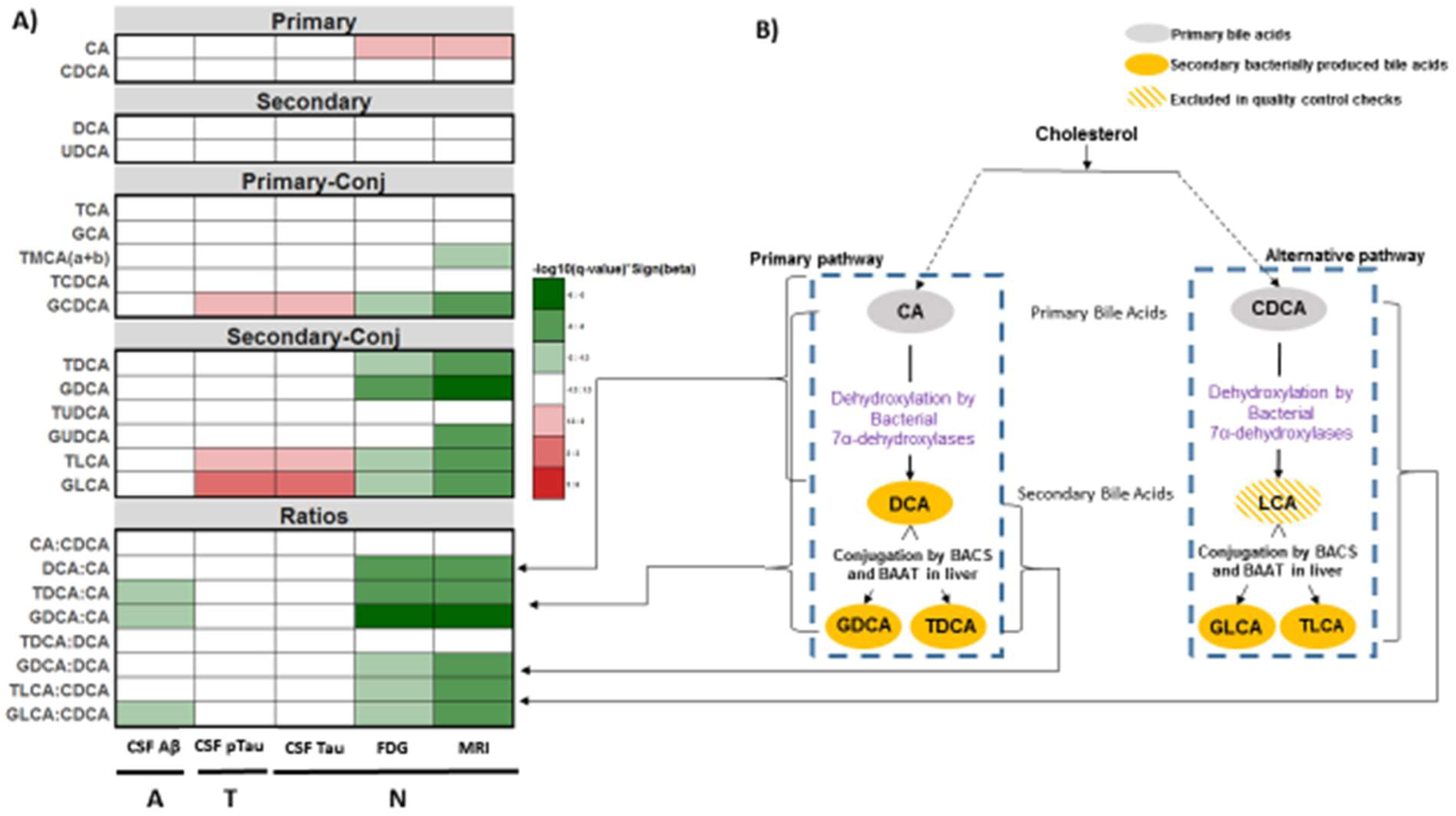
Bile acids and their ratios reflective of gut microbiome and liver enzymatic activities and their correlation with ATN biomarkers for Alzheimer’s disease. Heat map of q-values of association between bile acid profiles and the “A/T/N” biomarkers for AD (a). P-values estimated from linear regression analyses were corrected for multiple testing using FDR (q-value). Color code: white indicates q-value>0.05, reds indicate significant positive associations, and greens indicate significant negative associations. Several ratios were calculated to inform about possible enzymatic activity changes in AD (b). These ratios reflect: (1) Shift in bile acid metabolism from primary to alternative pathway, (2) Changes in gut microbiome correlated with production of secondary bile acids, and (3) Changes in glycine and taurine conjugation of secondary bile acids. LCA was excluded in prepossessing checks.

### 2.3. Neuroimaging processing

#### 2.3.1. Magnetic Resonance Imaging (MRI)

T_1_-weighted brain MRI scans at baseline were acquired using a sagittal 3D MP-RAGE sequence following the ADNI MRI protocol.[30, 31] As detailed in previous studies, FreeSurfer V5.1, a widely employed automated MRI analysis approach, was used to process MRI scans and extract whole brain and ROI (region of interest)-based neuroimaging endophenotypes including volumes and cortical thickness determined by automated segmentation and parcellation.[32-34] The cortical surface was reconstructed to measure thickness at each vertex. The cortical thickness was calculated by taking the Euclidean distance between the grey/white boundary and the grey/cerebrospinal fluid (CSF) boundary at each vertex on the surface.[35-37]

#### 2.3.2. Positron Emission Tomography (PET)

Pre-processed [^18^F] FDG and [^18^F] Florbetapir PET scans (co-registered, averaged, standardized image and voxel size, uniform resolution) were downloaded from the ADNI LONI site (http://adni.loni.usc.edu) as described in previously reported methods for acquisition and processing of PET scans from the ADNI sample.[32, 38] For [^18^F] FDG PET, scans were intensity-normalized using a pons ROI to create [^18^F] FDG standardized uptake value ratio (SUVR) images. For [^18^F] Florbetapir PET, scans were intensity-normalized using a whole cerebellum reference region to create SUVR images.

### 2.4. CSF Aβ1-42, t-tau, and p-tau181 biomarkers

ADNI generated CSF biomarkers (Aβ1-42, t-tau and p-tau181) in pristine aliquots of 2,401 ADNI CSF samples using the validated and highly automated Roche Elecsys^®^ electrochemiluminescence immunoassays[39, 40] and the same reagent lot for each of these three biomarkers. The CSF biomarker data was downloaded from the ADNI LONI site (http://adni.loni.usc.edu).

### 2.5. Statistical analyses

#### 2.5.1. CSF biomarkers

We performed a linear regression analysis using age, sex, study phase (ADNI-1 or ADNI-GO/2), body mass index (BMI), and *APOE* ε4 status as covariates, followed by false discovery rate (FDR)-based multiple comparison adjustment with the Benjamini-Hochberg procedure.

#### 2.5.2. Region of Interest (ROI) based analysis of structural MRI and PET

Mean hippocampal volume was used as an MRI-related phenotype. For FDG PET, a mean SUVR value was extracted from a global cortical ROI representing regions where AD patients show decreased glucose metabolism relative to cognitively normal older participants (CN) from the full ADNI-1 cohort, normalized to pons.[38] For [^18^F] Florbetapir PET, a mean SUVR value was extracted using MarsBaR from a global cortical region generated from an independent comparison of ADNI-1 [11C]Pittsburgh Compound B SUVR scans (regions where AD > CN). We performed a linear regression analysis using age, sex, BMI, study phase (ADNI-1 or ADNI-GO/2), and *APOE* ε4 status as covariates. For hippocampal volume, years of education, intracranial volume (ICV), and magnetic field strength were added as additional covariates. FDR-based multiple comparison adjustment with the Benjamini-Hochberg procedure was used because the AD biomarker phenotypes were strongly correlated with each other.[41] Not accounting for this high collinearity of dependent variables would lead to an overly stringent correction for multiple testing.

#### 2.5.3. Unbiased while brain imaging analysis

The SurfStat software package (www.math.mcgill.ca/keith/surfstat/) was used to perform a multivariate analysis of cortical thickness to examine the effect of BA profiles on brain structural changes on a vertex-by-vertex basis using a general linear model (GLM) approach.[37] GLMs were developed using age, sex, years of education, ICV, BMI, *APOE* ε4 status, and magnetic field strength as covariates. The processed FDG PET images were used to perform a voxel-wise statistical analysis of the effect of BA levels on brain glucose metabolism across the whole brain using SPM8 (www.fil.ion.ucl.ac.uk/spm/). We performed a multivariate regression analysis using age, sex, BMI, *APOE* ε4 status, and study phase (ADNI-1 or ADNI-GO/2) as covariates. In the whole brain surface-based analysis, the adjustment for multiple comparisons was performed using the random field theory correction method with p<0.05 adjusted as the level for significance.[42-44]. In the voxel wise whole brain analysis, the significant statistical parameters were selected to correspond to a threshold of *p* < 0.05 (FDR-corrected).

## 3. Results

### 3.1. Study sample after QC

After QC procedures, 1,562 ADNI participants with 23 bile acids and their relevant ratio levels at baseline (370 cognitively normal older adults (CN), 98 significant memory concern (SMC), 284 early MCI (EMCI), 505 late MCI (LMCI), and 305 AD) were available for analysis. Demographic information for the study population is presented in **Table 1**.

**Table 1.** Demographics of ADNI participants stratified by baseline diagnosis^a^.

| Variable | N | CN<br>(N=370) | SMC<br>(N=98) | EMCI<br>(N=284) | LMCI<br>(N=505) | AD<br>(N=305) |
| --- | --- | --- | --- | --- | --- | --- |
| Age | 1562 | 74.58(5.71) | 72.18(5.63) | 71.12(7.51) | 73.95(7.59) | 74.70(7.79) |
| Sex: Female, No. (%) | 1562 | 190(51) | 56(57) | 130(46) | 139(39) | 139(46) |
| Education, years | 1562 | 16.28(2.92) | 16.71(2.56) | 15.95(2.66) | 15.87(2.90) | 15.16(3.00) |
| BMI (Kg/M<sup>2</sup>) | 1562 | 27.05(4.46) | 28.22(6.24) | 28.06(5.41) | 26.54(4.25) | 25.83(4.69) |
| APOE ε4 status, (+%) | 1562 | 104(28) | 32(33) | 121(43) | 273(54) | 202(66) |

<sup>a</sup>Data are reported as mean(SD) unless otherwise indicated
*Abbreviations:* AD: Alzheimer's disease; BMI: Body mass index; CN: Cognitively normal; EMCI: Early mild cognitive impairment; LMCI: Late mild cognitive impairment; SMC: subjective memory complaint

### 3.2. Biomarkers of amyloid-β (“A”)

We used CSF Aβ1-42 levels and a global cortical amyloid deposition of amyloid PET as biomarkers of amyloid-β. First, we evaluated whether BA profiles were associated with CSF Aβ1-42 biomarker by performing an association analysis for 15 BA metabolites and 8 relevant ratios with *APOE* ε4 status as a covariate. As shown in **Fig 1a**, after applying FDR-based multiple comparison correction, we identified three BA ratios significantly associated with CSF Aβ1-42 levels. Regression coefficients of the three BA ratios of bacterially produced conjugated secondary BAs to primary BAs (GDCA:CA, TDCA:CA, and GLCA:CDCA) showed negative associations indicating higher levels were associated with lower CSF Aβ1-42 values (CSF Aβ1-42 positivity). However, global cortical amyloid deposition of amyloid PET was not significantly associated with any BA or their ratios after applying FDR-based multiple comparison correction. GDCA:CA was marginally associated with a global cortical amyloid load (uncorrected *p*-value < 0.05). Higher GDCA:CA levels were associated with greater amyloid deposition.

### 3.3. Biomarkers of fibrillary tau (“T”)

We used CSF phosphorylated tau (p-tau) levels as the biomarker of fibrillary tau. We investigated the association of 23 BAs and their relevant ratios with CSF p-tau, with *APOE* ε4 status included as a covariate. We identified three significant associations (FDR-corrected *p*<0.05) (**Fig 1a**). For one conjugated primary BA metabolite (GCDCA), higher GCDCA levels were associated with higher CSF p-tau values. For two bacterially produced conjugated secondary BA metabolites (GLCA and TLCA), higher levels were correlated with higher CSF p-tau values.

### 3.4. Biomarkers of neurodegeneration or neuronal injury (“N”)

We used atrophy on T1-weighted MRI, hypometabolism on FDG PET, and CSF total tau (t-tau) levels as biomarkers of neurodegeneration or neuronal injury.

#### 3.4.1. Structural MRI (atrophy)

We investigated the association of BA metabolites and ratios with mean hippocampal volume with *APOE* ε4 status as a covariate. Among 23 BA characteristics, 14 BAs/ratios were significantly associated with hippocampal volume after controlling for multiple testing using FDR (**Fig 1a**; corrected *p*<0.05). For one primary BA metabolite, lower CA levels were associated with decreased hippocampal volume. However, for two conjugated primary BA metabolites (GCDCA and TMCA(a+b)) and five bacterially produced conjugated secondary BA metabolites (GDCA, GLCA, GUDCA, TDCA, and TLCA), higher BA levels were associated with decreased hippocampal volume. In addition, higher levels of six ratios of bacterially produced secondary BA metabolite to primary BA metabolite (DCA:CA, GDCA:CA, TDCA:CA, GDCA:DCA, GLCA:CDCA, and TLCA:CDCA) were associated with decreased hippocampal volume.

Among the 14 significant BA signatures, six BA profiles were significantly associated with CSF Aβ1-42 biomarker (“A”) or CSF p-tau biomarker (“T”). For the six BA profiles, we performed a detailed whole-brain surface-based analysis using multivariate regression models and assessed their effects on whole-brain cortical thickness in an unbiased way. We identified significant associations for all six BA profiles (cluster wise threshold of RFT-corrected *p* < 0.05), which showed consistent patterns in the associations of CSF Aβ1-42 or p-tau levels (**Fig 2**). Higher levels of a conjugated primary BA (GCDCA) were significantly associated with reduced cortical thickness especially in bilateral entorhinal cortices. Increased levels of one bacterially produced conjugated secondary BA metabolite (GLCA) and two ratios of bacterially produced secondary BA metabolites to primary BA metabolites (GDCA:CA and GLCA:CDCA) were significantly associated with reduced cortical thickness in the bilateral frontal, parietal, and temporal lobes including the entorhinal cortex. For one bacterially produced conjugated secondary BA metabolite (TLCA) and one ratio of a bacterially produced secondary BA metabolite to a primary BA metabolite (TDCA:CA), increased levels were associated with reduced cortical thickness in a widespread pattern, especially in the bilateral frontal, parietal, and temporal lobes.

**Fig 2.**
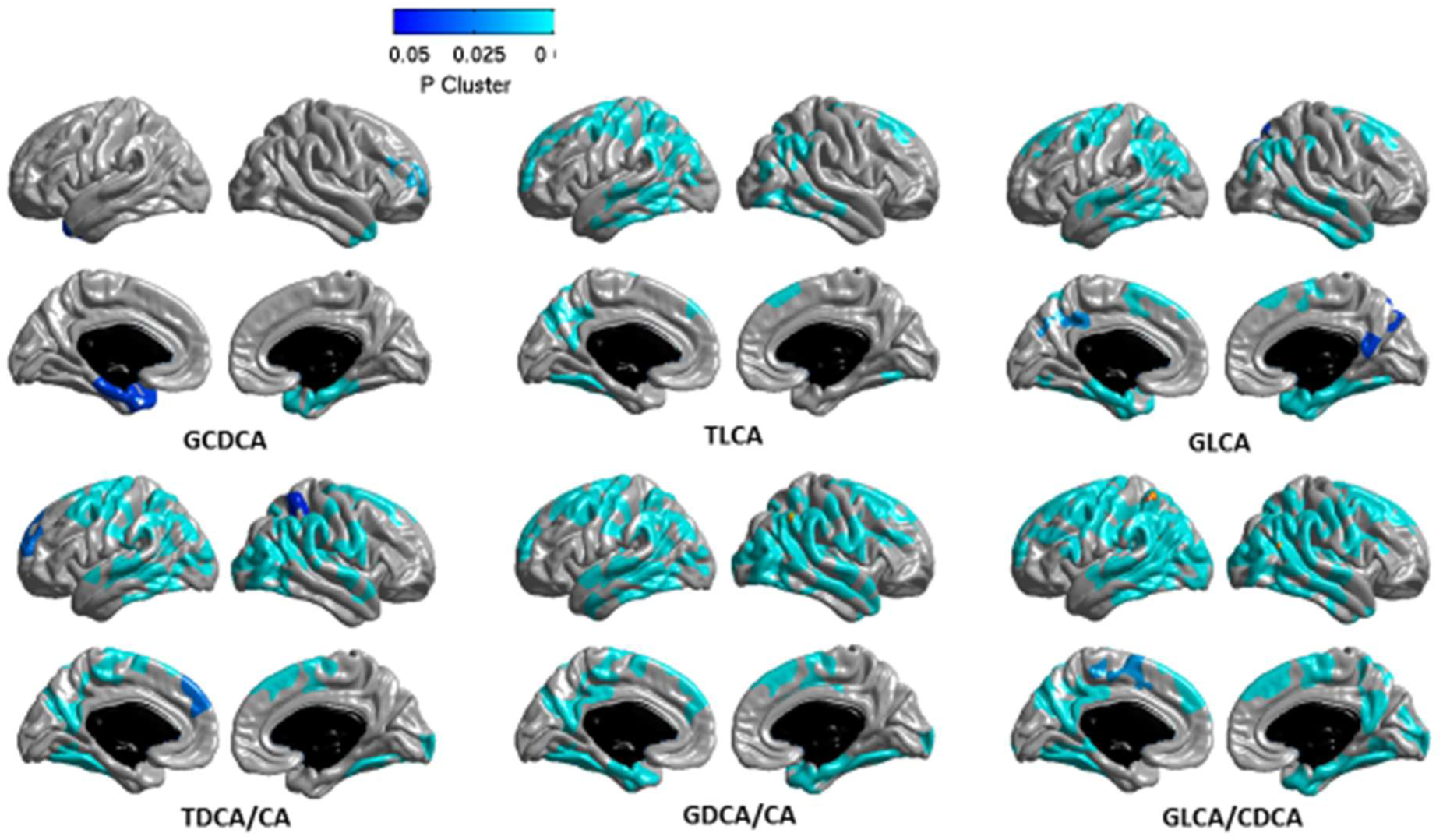
Unbiased whole-brain surface-based imaging analysis. A whole-brain multivariate analysis of cortical thickness across the brain surface was performed to visualize the topography of the association of bile acid profiles with brain structure in an unbiased manner. For a surface-based analysis of levels of CDCA, TLCA, GLCA, TDCA:CA, GDCA:CA, and GLCA:CDCA, statistical maps were thresholded using a random field theory for a multiple testing adjustment to a corrected significance level of 0.05. The *p*-value for clusters indicates significant corrected *p* values with the lightest blue color. Higher GCDCA levels were significantly associated with reduced cortical thickness especially in bilateral entorhinal cortices. Increased GLCA, GDCA:CA, and GLCA:CDCA levels were significantly associated with reduced cortical thickness in the bilateral frontal, parietal, and temporal lobes including the entorhinal cortex. For TLCA and TDCA:CA, increased levels were associated with reduced cortical thickness in a widespread pattern, especially in the bilateral frontal, parietal, and temporal lobes.

#### 3.4.2. FDG-PET (brain glucose metabolism)

We performed an association analysis for 23 BA and ratios with global cortical glucose metabolism measured by FDG PET scans across 1,066 participants with both FDG PET scans and BA measurements. The association testing including *APOE* ε4 status as a covariate, identified twelve BA characteristics as significantly associated with brain glucose metabolism after controlling for multiple testing using FDR (**Fig 1a**; corrected *p*<0.05). For one primary BA metabolite, lower CA levels were associated with reduced glucose metabolism. In contrast, for one conjugated primary BA metabolite (GCDCA), four bacterially produced conjugated secondary BA metabolites (GDCA, GLCA, TDCA, and TLCA), and six ratios of bacterially produced secondary BA metabolites to primary BA metabolites (DCA:CA, GDCA:CA, TDCA:CA, GDCA:DCA, GLCA:CDCA, and TLCA:CDCA), higher BA ratio levels were associated with reduced glucose metabolism.

In addition, in an unbiased way we performed a detailed whole-brain analysis to determine the effect of BAs on brain glucose metabolism on a voxel wise level for six BAs and ratios (GCDCA, GLCA, TLCA, GDCA:CA, TDCA:CA, and GLCA:CDCA) that were significantly associated with both CSF Aβ1-42 or p-tau biomarkers, FDG metabolism, and hippocampal volume. We identified significant associations for all six BA profiles (cluster wise threshold of FDR-corrected *p* < 0.05), which showed consistent patterns in the associations of CSF Aβ1-42 or p-tau levels and structural atrophy (**Fig 3**). Higher levels of a conjugated primary bile acid GCDCA were significantly associated with reduced glucose metabolism especially in the bilateral hippocampi, which showed consistent patterns with the associations of cortical thickness. Increased levels of one bacterially produced conjugated secondary BA metabolite (GLCA) and one ratio of a bacterially produced secondary BA metabolite to a primary BA metabolite (GLCA:CDCA) were significantly associated with reduced glucose metabolism in the bilateral temporal and parietal lobes. Lower TLCA levels, a bacterially produced conjugated secondary BA metabolite, were associated with increased glucose metabolism in the left temporal lobe. For two ratios (GDCA:CA and TDCA:CA) of bacterially produced secondary BA metabolite to a primary BA metabolite, higher ratio levels were significantly associated with reduced glucose metabolism in a widespread pattern, especially in the bilateral frontal, parietal, and temporal lobes.

**Fig 3.**
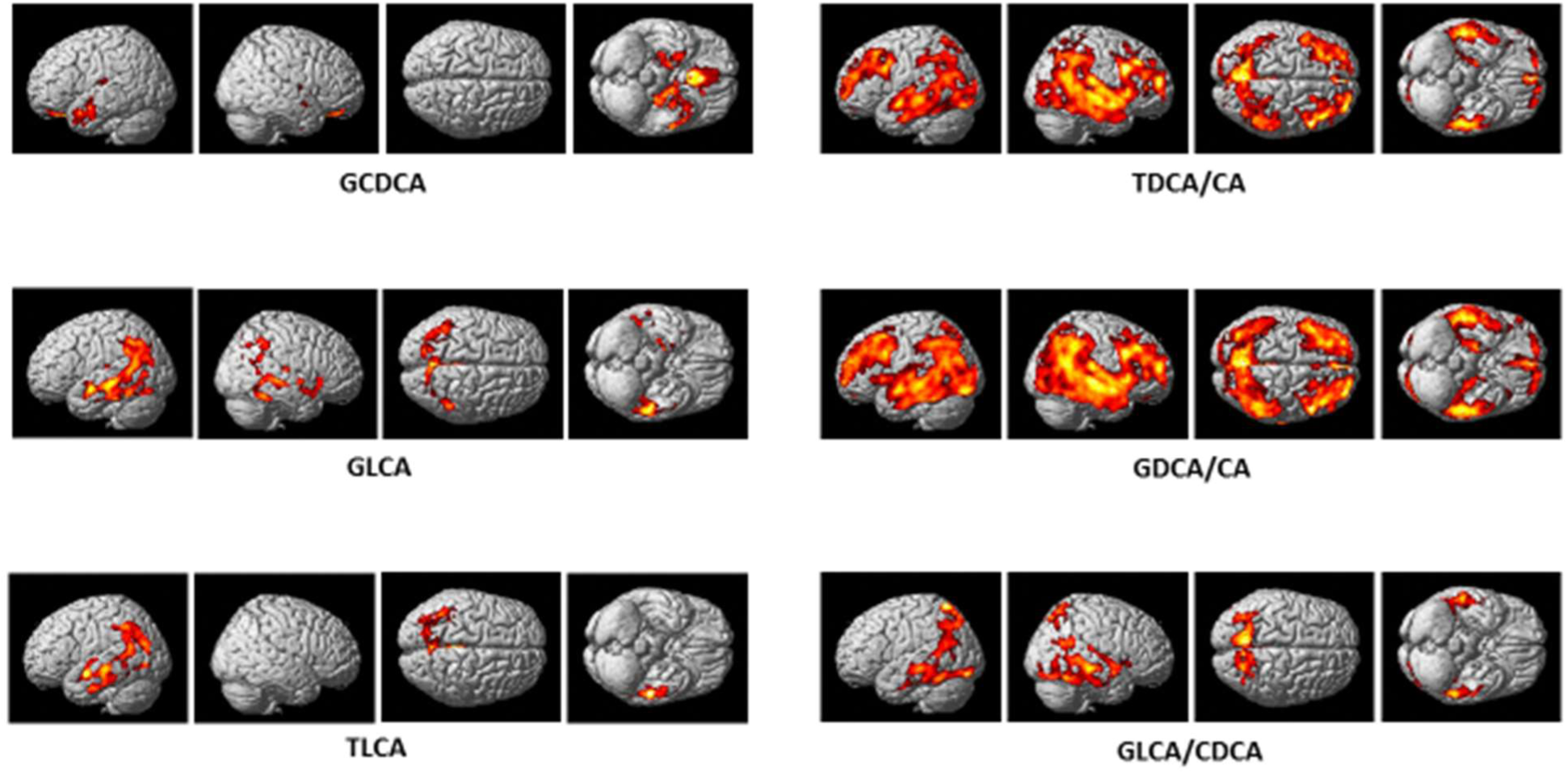
Unbiased whole-brain voxel-based imaging analysis. A whole-brain multivariate analysis of glucose metabolism was performed to visualize the topography of the association of bile acid profiles with glucose metabolism in an unbiased manner. For a voxel-based analysis of FDG-PET scans, we identified significant associations (cluster wise threshold of FDR-corrected *p* < 0.05). Higher GCDCA levels were significantly associated with reduced glucose metabolism especially in the bilateral hippocampi. Increased GLCA and GLCA:CDCA levels were significantly associated with reduced glucose metabolism in the bilateral temporal and parietal lobes. Lower TLCA levels were associated with increased glucose metabolism in the left temporal lobe. For two ratios (GDCA:CA and TDCA:CA), higher ratio levels were significantly associated with reduced glucose metabolism in a widespread pattern, especially in the bilateral frontal, parietal, and temporal lobes.

#### 3.4.3. CSF total tau (t-tau)

We evaluated whether 23 BAs and ratios were associated with the CSF t-tau including *APOE* ε4 status as a covariate. We identified three significant associations after controlling for multiple testing using FDR (corrected *p*<0.05) (Fig. 1). Higher levels of GCDCA, a conjugated primary BA metabolite, and GLCA and TLCA, bacterially produced secondary BA metabolites, were associated with higher CSF t-tau values.

## 4. Discussion

In this report we analyzed serum-based BA profiles in the ADNI cohort to investigate the relationship between peripheral metabolic measures and central biomarkers for AD pathophysiology based on the recently proposed framework (“A/T/N”).[27] Our results showed that altered BA profiles were significantly associated with structural and functional changes in brain as noted by larger atrophy and reduced glucose metabolism in brain (“N”). Furthermore, altered BA profiles were significantly associated with three CSF biomarkers including Aβ1-42, t-tau, and p-tau. Three ratios of primary BAs to secondary BAs were associated with lower CSF Aβ1-42 levels (amyloid-β positivity) (“A”) as well as reduced cortical glucose metabolism and larger structural atrophy (GDCA:CA, TDCA:CA, and GLCA:CDCA). One conjugated primary BA profile (GCDCA) and two bacterially produced conjugated secondary BAs (GLCA and TLCA) were associated with higher CSF p-tau values (“T”) as well as higher CSF t-tau values, reduced glucose metabolism, and larger structural atrophy.

Whether the gut microbiome directly influences AD pathogenesis remains unknown, however is does appear to influence amyloid-β, fibrillary tau, and neurodegeneration. To our knowledge, this is the first study to systematically link markers of the gut microbiome and liver function to AD-related structural and functional neuroimaging biomarkers as well as biomarkers of amyloid-β and tau burden.

Three core CSF biomarkers (Aβ1-42, t-tau, and p-tau) reflect AD pathology and can be used to reliably diagnose AD and identify MCI, a prodromal stage of AD, with high diagnostic accuracy.[45, 46] Previous studies showed that AD patients have a substantial reduction in CSF Aβ1-42 and a marked increase in levels of CSF t-tau and p-tau[47-50]. We observed that higher levels of TDCA:CA, GDCA:CA, and GLCA:CDCA were associated with decreased levels of CSF Aβ1-42 and higher levels of GCDCA, TLCA, and GLCA were associated with increased levels of CSF t-tau and p-tau.

MRI is widely used to investigate structural changes in MCI and AD.[51-53] We observed lower levels of CA and higher levels of GCDCA, TMCA(a+b), GDCA, GLCA, GUDCA, TDCA, TLCA, DCA:CA, GDCA:CA, TDCA:CA, GDCA:DCA, GLCA:CDCA, and GCDCA:CDCA were associated with greater brain atrophy. Significant regional effects were observed particularly in the bilateral inferior parietal gyri cortices, hippocampi, and temporal lobes including the entorhinal cortex. The hippocampus and entorhinal cortex are affected early in AD, and the decrease in hippocampal volume accelerates as AD progresses. Significant thinning of the cortical surface reflects atrophy in the temporal, parietal, and frontal lobes has been shown in MCI and AD.[51-53] Reduced cortical thickness in the temporal cortex as a measure of brain atrophy rate has shown promise in predicting MCI to AD progression.[51]

Lower CA and higher TDCA, GDCA, GCDCA, GLCA, TLCA, DCA:CA, GDCA:CA, TDCA:CA, GDCA:DCA, GLCA:CDCA, and TLCA:CDCA levels were associated with reduced global glucose metabolism in brain. The significant regional effect on brain glucose metabolism was observed particularly in the bilateral hippocampi for GCDCA, in the temporal and parietal lobes for GLCA, TLCA, and GLCA/CDCA, and in a widespread pattern including the bilateral temporal, parietal, and frontal lobes for GDCA:CA and TDCA:CA. AD patients have shown significant glucose metabolism reduction in the temporal lobes, parietal lobes, and then the frontal lobes with increasing severity of AD.[54-56]

The observed pattern of association between changes in brain structure and glucose metabolism as well as CSF biomarkers with specific BAs and ratios indicates a potential mechanistic connection between peripheral and central biochemical changes. Our results strongly suggest gut-liver-brain axis involvement in AD, neurodegeneration, and brain dysfunction. Both liver function and gut microbiome activity are impacted in AD, and these changes seem to occur at the earliest stages of disease. Despite this strong pattern of associations, the specific mechanism and causal directionality remains to be determined.

We hypothesized that altered gut microbiota play an important role. This is supported by several lines of research connecting the gut microbiota and AD pathology. Alterations in the gut microbiota and an increase in gut permeability may lead to dysfunction in the hippocampus[57, 58] and the development of insulin resistance, which correlates with AD pathogenesis.[59-61] It has been hypothesized that increased gut permeability allows bacteria-derived amyloids from the gastrointestinal tract to accumulate at the systemic and brain level[62]. This in turn could lead to the upregulation of pro-inflammatory microRNA-34a and as a consequence, downregulation of TREM2, leading to the accumulation of Aβ_42._[61, 62]

Results from animal studies demonstrate that increased input of BAs significantly inhibits two of the major phyla in the human gut microbiome, Bacteroidetes and Actinobacteria.[63] Bacterial taxonomic composition of fecal samples revealed differences in bacterial abundance including decreased Firicutes, increased *Bacteroidetes*, and decreased *Bifidobacterium* (phylum Actinobaceria) in the microbiome of AD patients relative to age- and sex-matched controls. Furthermore, these same differences in bacterial abundance correlated with CSF biomarkers including Aβ_42_/Aβ_40_ and p-tau/Aβ_42_. Even in the age- and sex-matched controls (no dementia diagnosis), there was a similar relationship between the same bacteria that were either more or less abundant in AD and markers of tau and amyloid.[64] In another study, an increased abundance of pro-inflammatory bacteria (*Escherichia/Shigella)* and a decreased abundance of anti-inflammatory bacteria (*Eubacterium* rectale) were noted in cognitively impaired older adults with evidence of amyloid deposition on PET imaging compared to those who were amyloid negative.[65] These results lend further support to the link between gut microbiota and brain amyloidosis. Gut microbiota have been associated with the accumulation of amyloid plaques in a mouse model of AD. A transgenic AD mouse model generated under germ-free conditions had dramatic reductions in cerebral amyloid-β pathology compared to control animals with normal intestinal microbiota. More intriguingly, colonization of germ-free AD mice with microbiota harvested from conventionally raised AD mice showed dramatic increases in Aβ pathology.[25]

Decreasing BA in the gut favors gram-negative members of the microbiome, leading to the production of lipopolysaccharides (LPS) which elicits a strong immune response. Furthermore, overgrowth of gram-negative bacteria could promote insulin resistance.[57] Lipopolysaccharides concentrations are elevated in AD when compared to controls.[66, 67] In a mouse model of AD, peripheral inflammation induced through LPS resulted in significantly higher levels of Aβ1-42 in the hippocampus and marked cognitive deficits[68]. Several studies have confirmed that increased inflammation as a result of LPS is associated with memory impairments.[69-71] It is thought that declines in memory are the result of the association between LPS-accumulation of Aβ and neuronal cell death.[69] In contrast, increased BA in the gut favors gram-positive members of Firmictures, including bacteria that 7α-dehydroxylate host primary BAs into secondary toxic BAs.[72-74]

The association between BA cytotoxicity and the generation of reactive oxygen species (ROS) is well documented.[75-79] Others have proposed that mitochondrial ROS production plays an important role in brain metabolic signaling.[80, 81] Some of the mechanisms by which mitochondrial dysfunction leads to neuronal degeneration in AD include ROS generation and activation of mitochondrial permeability transition[82, 83], suggesting a crucial role for oxidative stress in the pathophysiology of AD. Hyperphosphorylation of tau proteins has been linked to oxidation through the microtubule-associated protein kinase pathway.[84] In our analyses, three cytotoxic BAs (GCDCA, GLCA, and TLCA) correlated with higher biomarker levels of fibrillary tau and neurodegeneration/neuronal injury.

Hydrophobic BAs, like CDCA, are known to damage biological membranes[85], whereas hydrophilic BAs, like UDCA and TUDC, are inhibitors of apoptosis via their ability to stabilize mitochondrial membranes.[86, 87] Impairment of mitochondrial function is likely one of the vital ways in which BAs cause cellular dysfunctions[88-91]. Decreased mitochondrial membrane potential has been associated with increasing concentrations of the bile acids, LCA, DCA, UDCA, CDCA, GCDC, and taurochenodeoxcholic (TCDC) acid.[88]

## 4.1. Limitations

The ADNI study is observational by design making it difficult to control for confounding as well as to determine directionality of associations and causal pathways. For example, the population of the gut is affected by a plethora of factors including geography, lifetime immunological experience, and environmental factors, which could play important, yet currently unknown roles in the pathogenesis of AD. Experimental studies are needed to understand the mechanistic role of BA in the development of AD-related pathology as well as to disentangle cause and effect. Medication use was extensively explored as a potential confounder (**Supplementary Fig 1**). Overall, our key findings remained significant after adjustment for medication use, although effects were attenuated. In addition, further studies are warranted to validate our findings in independent cohorts.

## 5. Conclusions

This is the first study to our knowledge to demonstrate an association between altered BA profiles and amyloid-β, tau, and neurodegeneration biomarkers of AD pathophysiology. While our results provide further evidence implicating BA signaling in AD, the causal pathway remains to be systematically investigated by prospective clinical studies and experimental manipulations in model systems. Future metagenomics studies are also needed to define the relationship between BAs, host factors including genetics, and bacterial community composition within an individual across time. Building on present results, these investigations are needed to achieve a mechanistic understanding of the role of gut bacteria and BAs in relation to AD pathophysiology. If a causal role can be demonstrated in future research, BA signaling pathways may lead to the identification of metabolites that are protective against AD and could foster novel therapeutic strategies.

## Author Contributions

Nho, MahmoudianDehkordi, Arnold had full access to all of the data in the study and take responsibility for the integrity of the data and the accuracy of the data analysis.

**Statistical analyses also included:** Arnold, Kastenmüller

**Data management and medication term mapping:** Blach

**Concept and design:** Kaddurah-Daouk led concept and design team that included all co-authors

**Drafting of the manuscript:** Nho, Kueider-Paisley, Saykin, Kaddurah-Daouk

**Biochemical, genomics and medications integration:** Kastenmüller, Baillie, Han, Risacher

**Data deposition:** Alzheimer’s Disease Neuroimaging Initiative (see note)

**Harmonization of methods:** Alzheimer’s Disease Metabolomics Consortium (see note)

**Technical, bibliographic research and/or material support:** Louie

**Biochemical interpretation:** Baillie, Han, Kaddurah-Daouk

**Critical revision of the manuscript for important intellectual content:** Saykin, Doraiswamy, Kaddurah-Daouk

**Obtained funding:** Kaddurah-Daouk

**Supervision:** Trojanowski, Shaw, Weiner, Doraiswamy, Saykin, Kastenmüller, Kaddurah-Daouk

## The Alzheimer’s Disease Neuroimaging Initiative (ADNI)

Data used in the preparation of this article were obtained from the ADNI database (http://adni.loni.usc.edu). As such, the investigators within the ADNI contributed to the design and implementation of ADNI and/or provided data but did not participate in analysis or writing of this report. A complete listing of ADNI investigators can be found at: http://adni.loni.usc.edu/wp-content/uploads/how_to_apply/ADNI_Acknowledgement_List.pdf.

## Additional Contributions

The authors are grateful to Lisa Howerton for administrative support and the numerous ADNI study volunteers and their families.

## DISCLOSURES

J.Q.T. may accrue revenue in the future on patents submitted by the University of Pennsylvania wherein he is a co-inventor and he received revenue from the sale of Avid to Eli Lily as a co-inventor on imaging-related patents submitted by the University of Pennsylvania. L.M.S. receives research funding from NIH (U01 AG024904; R01 MH 098260; R01 AG 046171; 1RF AG 051550); MJFox Foundation for PD Research and is a consultant for Eli Lilly; Novartis; Roche; he provides QC over-sight for the Roche Elecsys immunoassay as part of responsibilities for the ADNI study. A.J.S. reports investigator-initiated research support from Eli Lilly unrelated to the work re-ported here. He has received consulting fees and travel expenses from Eli Lilly and Siemens Healthcare and is a consultant to Arkley BioTek. He also receives support from Springer publishing as an editor in chief of Brain Imaging and Behavior. M.W.W. reports stock/stock options from Elan, Synarc, travel expenses from Novartis, Tohoku University, Fundacio Ace, Travel eDreams, MCI Group, NSAS, Danone Trading, ANT Congress, NeuroVigil, CHRU-Hopital Roger Salengro, Siemens, AstraZeneca, Geneva University Hospitals, Lilly, University of California, San Diego–ADNI, Paris University, Institut Catala de Neurociencies Aplicades, University of New Mexico School of Medicine, Ipsen, Clinical Trials on Alzheimer’s Disease, Pfizer, AD PD meeting. PMD has received research grants and advisory/speaking fees from several companies for other projects, and he owns stock in several companies. Full disclosures will be made through the IJCME form. R.K.D. is inventor on key patents in the field of metabolomics including applications for Alzheimer disease. All other authors report no disclosures.

An email with links to the Authorship Form will be sent to authors for completion after manuscripts have been submitted

